# From Nagelkerke’s R^2^ to Liability-Scale Variance Explained for Polygenic Scores

**DOI:** 10.64898/2026.09.12.751172

**Authors:** Emil Uffelmann, Peter M. Visscher

**Author notes:** Address correspondence to Emil Uffelmann; or Peter Visscher.

## Abstract

It is desirable to quantify the prediction accuracy of polygenic scores (PGS) for disease on the scale of liability and adjusted for case-control ascertainment in the test sample, because that allows comparison across prevalence and ascertainment. Previous expressions have focused in their derivation and implementation on linear regression on the observed 0-1 scale followed by a transformation of the coefficient of determination (*R*^2^) to the scale of liability, adjusted for ascertainment. Yet most statistical analyses with empirical data use logistic regression. The differences in scale have led to confusion and incorrect transformations in the literature. Here we provide a new derivation and simple equation, validated by simulation, that allows a direct transformation from the empirical results from logistic regression to the scale of liability.

## Introduction

Coefficients of determination (*R*^2^) quantify the proportion of phenotypic variance explained by polygenic scores (PGSs) and are the standard metric for reporting their predictive accuracy. For continuous phenotypes, *R*^2^ is equal to the squared correlation between the PGS and the outcome. For binary disease phenotypes, *R*^2^ computed from a linear regression on the observed 0-1 scale 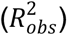 depends on both the population prevalence of the disease and the proportion of cases in the test sample, so values are not comparable across diseases with differing prevalences or across studies with differing ascertainment.

Under the liability threshold model (Dempster & Lerner, 1950; Lynch & Walsh, 1998), disease status is determined by an unobserved, normally distributed liability, and individuals are affected when their liability exceeds a threshold determined by the population prevalence. This model can be utilized to compare diseases on a continuous scale unaffected by prevalence or ascertainment. Following this reasoning, Lee and colleagues derived a widely used conversion from 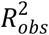(i.e., the binary disease scale) to the unobserved continuous liability scale 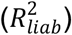 (Lee et al., 2012), adapting a transformation introduced earlier for SNP-based heritability (Golan et al., 2014; Lee et al., 2011).

In practice, binary phenotypes tend to be analyzed using logistic regression, which yields no measure directly analogous to the *R*^2^ in linear regression. Instead, several pseudo- *R*^2^ metrics have been developed, of which Nagelkerke-*R*^2^ 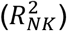 (Nagelkerke, 1991) is arguably the most popular. Like 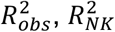 on both the population prevalence and the proportion of cases in the sample (Choi et al., 2020), and so cannot be compared across studies or diseases. 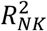 is sometimes passed through the Lee conversion as though it were 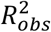(Lewis & Vassos, 2020; Nicolas et al., 2025), and the accuracy of the resulting liability-scale estimates has not, to our knowledge, been evaluated.

Here we present a new derivation that leads to a simple adaptation of the Lee conversion that takes 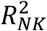 as input in place of 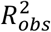. Using simulations with different population prevalences and ground truth 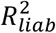 values, we show that the adapted conversion recovers the true 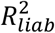 across realistic values, whereas the original Lee conversion applied to 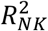 is systematically upward biased.

## Results

### Converting 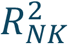to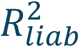

We derive the conversion from the liability threshold model, using only the population prevalence (*K*) and the test sample case fraction *P* = Pr(*Y* = 1), with *Y* ∈ {0,1} denoting disease status. We consider logistic regression with a single covariate *X* so that the probability of disease is 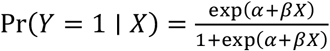. Without loss of generality, *X* is centered so that E(*X*) = 0. The Cox and Snell’s *R*^2^ (Cox & Snell, 1989) is defined as 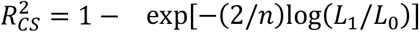, with *n* the sample size, *L*_1_ and *L*_O_ the likelihoods for the full model (fitting *α* and *β*) and null model (fitting *α* only), respectively. For a large sample size and small value of *β*, 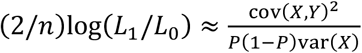 (see Appendix A). Therefore,

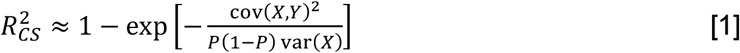

Since *Y* is binary, this can also be expressed as,

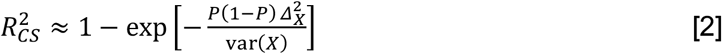

with *Δ*_*X*_ the difference in the mean covariate *X* between cases and controls in the test sample; 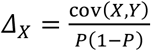. If we further assume that 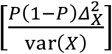is small, then

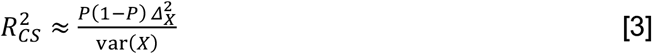

Note that *P*(1 − *P*) = var(*Y*). Therefore, 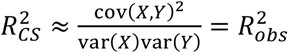, the coefficient of determination on the observed 0-1 scale in the test sample (Lee et al., 2012). Equation [3] is an approximation for any covariate. Nagelkerke’s *R*^2^ value is a simple scaled version of the Cox-Snell statistic, 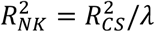, with *λ* = 1 − *P*^2*P*^(1 − *P*)^2(1–*P*)^.

For the special case of a PGS (*X* = *g*), we need to substitute the expected values of the mean difference in the PGS between cases and controls and the sample variance for *Δ*_*X*_ and var(*X*), respectively. These are known from the literature for a liability threshold model (Golan et al., 2014; Lee et al., 2011, 2012; Lynch & Walsh, 1998). In particular, for *t* = *Φ*^−1^(1 − *K*) the liability threshold, *z* = *ϕ*(*t*) the height of the standard normal density curve at threshold *t*, and 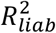 the variance in liability explained by the PGS in the population,

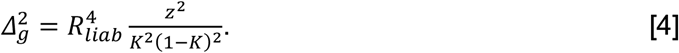

The variance of the PGS in an ascertained sample is

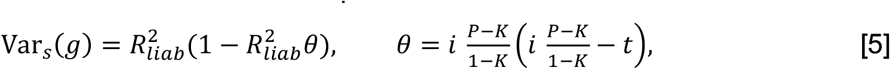

as previously derived in (Daetwyler et al., 2008; Golan et al., 2014; Lee et al., 2011). Substituting the threshold-model expressions into 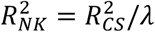 gives

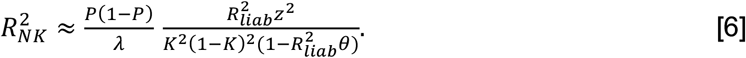

defining

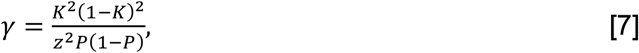

this simplifies to

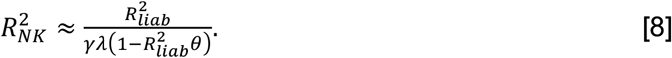

Solving for 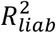 yields

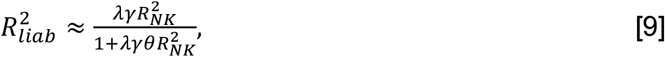

which is a simple adjusted version of the conversion from the observed linear scale to the liability scale, introduced in (Lee et al., 2012),

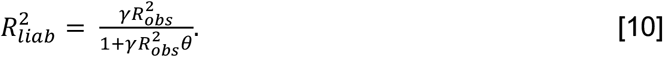

### Simulations

We evaluated the calibration of the new conversion in simulations based on the liability threshold model. We simulated individual-level data for 1000 SNPs in linkage equilibrium, of which 500 were causal, with a SNP-based heritability on the liability scale of 1 (a PGS cannot explain more liability variance than the total narrow-sense heritability, so the heritability was set to 1 to allow the target 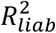 to span the full range up to 0.9). For each parameter setting, we simulated a discovery sample to estimate SNP effects using a case-control GWAS, and a testing sample (*N*_*case*_ = 1000, *N*_*control*_ = 1000) in which the PGS was computed and evaluated. The discovery sample size was set with the avengeme package (Dudbridge, 2013) to achieve a target 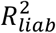 of the PGS in the testing sample, and we varied this target over 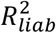 ={0.1, 0.2, 0.3, 0.4, 0.5, 0.6, 0.7, 0.8, 0.9} and the population lifetime prevalence over K = {0.01, 0.15}. Both the discovery and testing samples were ascertained at a 50% case-control ratio. We repeated the simulations 100 times for every parameter setting.

In the testing sample, we estimated the 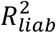 of the PGS in four ways. As a reference, we converted the 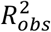to 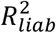 using the original Lee conversion (Lee et al., 2012) shown in equation [10]. We then estimated 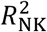 and similarly converted it using the original Lee equation [10] (the naive conversion) and our new equation [9] (the adjusted conversion).

The conversion based on equation [9] (i.e., adjusted conversion) was very well calibrated over most of the simulated range (see **Figure 1**). Only at low prevalence and unrealistically high 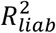did it become downward biased. In contrast, the naive conversion overestimated 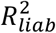at both prevalences, and the overestimation increased with increasing 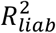. Reporting 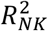 unconverted (no conversion) overestimated 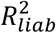at K = 0.01, but at K = 0.15 it turned into an underestimate above 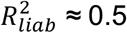(0.837 at a true value of 0.9). Both biases were most severe at the low prevalence, but peaked at different points of the range: at K = 0.01, 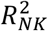(no conversion) was most inflated in the middle of the range (0.624 [95%-CI: 0.618–0.630] at a true 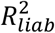of 0.4), whereas the naive conversion was most inflated at the top (1.683 [95%-CI: 1.672–1.693] at a true 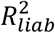 of 0.9). The naive conversion exceeded 1, an impossible value for a proportion of variance explained.

**Figure 1.**
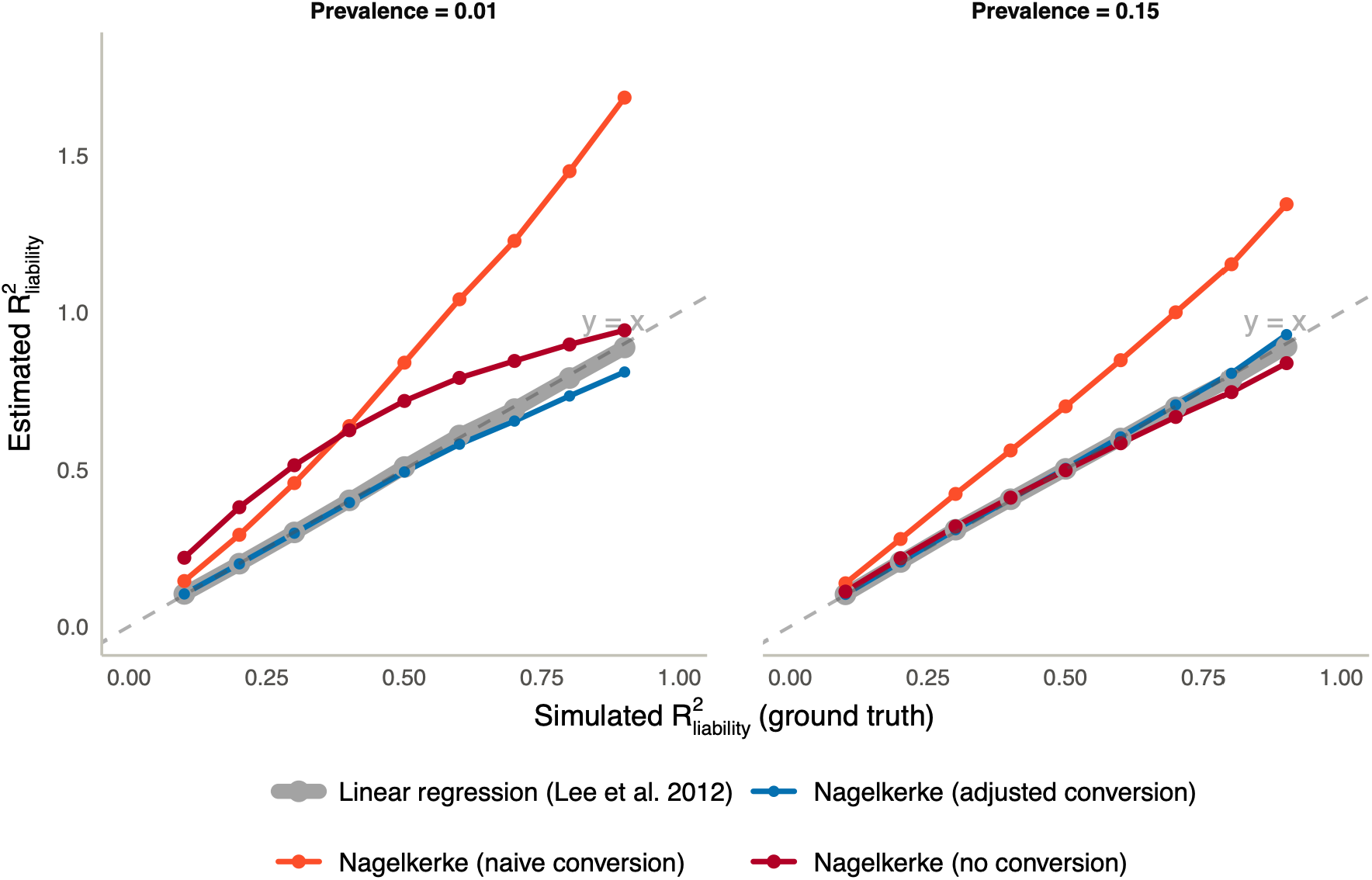
Conversions of *R*^2^ values to the liability scale. Estimated 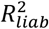 plotted against the simulated ground truth, for a population prevalence of *K* = 0.01 (left) and *K* = 0.15 (right). Each point is the mean across 100 replicates; the dashed line is the identity line (y = x). Applying the Lee conversion (equation [10]) to 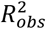 from linear regression, for which it is intended, and applying our conversion (equation [9]) to 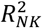 both recover the true 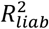 over most of the range. Applying the Lee conversion directly to 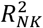 (naive conversion) overestimates 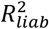 at both prevalences, while reporting 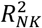 unconverted (no conversion) overestimates it at K = 0.01 and underestimates it at K = 0.15 above 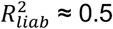.

We note that the Lee et al. transformation, based on linear regression of disease status in the case-control sample, is perfectly calibrated across the entire range of parameter values considered. This follows because, under the assumed liability-threshold model and ascertainment scheme, the means, variances, and covariances that determine the observed-scale 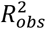 can be expressed exactly in terms of the corresponding population parameters.

## Discussion

Nagelkerke’s *R*^2^ 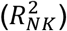is the most commonly reported pseudo-*R*^2^ for binary disease traits and is readily provided by software tools such as *glm*() in *R*, but is a function of both the population prevalence and the sample case fraction, preventing valid comparisons across diseases and studies. Variance explained on the liability scale 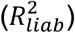 is independent of both, making it the preferred metric. Lee et al. (2012) introduced a canonical conversion for linear regression 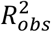 to 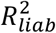, but it cannot be applied validly to 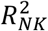. Nevertheless, this conversion has been incorrectly applied to 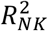 in the literature (Lewis & Vassos, 2020; Nicolas et al., 2025).

Here, we derived a simple conversion from 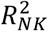 to 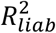 based on the liability threshold model, requiring only the population disease prevalence and the sample case fraction, and it is analogous to an adjusted version of the 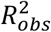to 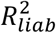conversion introduced in Lee et al. (2012). In simulations, our conversion was well calibrated across prevalences and realistic values of 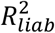, while a naive conversion of 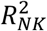 using the original Lee equation was not. Because the conversion requires only the population prevalence and the case fraction, previously published 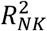 estimates can easily be converted retrospectively, allowing existing results to be placed on a comparable scale without reanalysis of individual-level data. To promote the application of the new conversion, we point to an *R* function which can be found on GitHub (see https://github.com/kn3in/ABC/blob/master/functions.R) and is reproduced in Appendix B.

We note that the conversion assumes a single predictor in the logistic regression model. Because other covariates that are associated with a disease of interest can become correlated with the PGS in an ascertained sample due to collider bias (Choi et al., 2020; Pirinen et al., 2012), correcting for these covariates can deflate 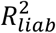 and therefore give biased results.

In summary, we derived a simple conversion of 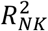 to 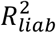that depends only on the population prevalence and the test sample case fraction, which was well calibrated in simulations across realistic values of 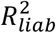 and can be applied easily in practice.

## Acknowledgements

P.M.V. acknowledges funding from the Australian Research Council (FL180100072) and the European Research Council (grant agreement no. 101198904), and E.U. acknowledges funding from the Netherlands Organization for Scientific Research - Gravitation project ‘BRAINSCAPES: A Roadmap from Neurogenetics to Neurobiology’ (024.004.012). During the preparation of this manuscript, the authors used Claude (Anthropic) and ChatGPT (OpenAI) to review the writing and derivations and to help design the figure. The authors reviewed and edited all content and take full responsibility for it.

## Conflict of interest

P.M.V. is an advisor to Starling Genomics and holds an indirect financial interest in the company.

## Data availability statement

This study used no empirical data; all analyses are based on simulated data generated as described in the manuscript. Appendix B provides the R function implementing the conversion.

## Appendix A

Cox and Snell’s *R*^2^ (Cox & Snell, 1989) is defined through the logistic regression likelihood ratio,

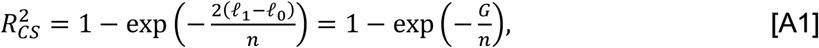

where *l*_0_ and *l*_1_ are the null and fitted log-likelihoods and *G* = 2(*l*_1_ – *l*_0_) is the likelihood-ratio statistic. For 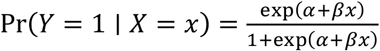, the score (*S*) and Fisher information (*I*) for *β* are *n* cov(*X, Y*) and *n* var(*Y*)var(*X*), respectively. A second-order expansion of 2(*l*_1_ − *l*_O_) around 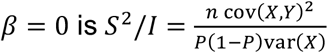, and therefore

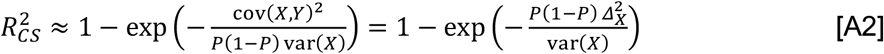

with *Δ*_*X*_ = E(*X* ∣ *Y* = 1) − E(*X* ∣ *Y* = 0).

## Appendix B

~~~
prs_r2nagelkerke_to_r2liab <- function(K,P,prs_r2nk){
  ## Adapted from Lee et al. 2012 Genet Epidemiology
  t = -qnorm(K,mean=0,sd=1) # disease threshold
  z<-dnorm(t)        # height of the normal distribution at T
  i1<-z/K            # mean liability of A1 (eg Falconer and Mackay)

  theta = i1*(P-K)/(1-K)*(i1*(P-K)/(1-K)-t)          # theta, eq. (15) in Lee et al. 2012
  cv = K*(1-K)/z^2*K*(1-K)/(P*(1-P))    # C in equation (15)
  lambda = 1 - P^(2*P) * (1-P)^(2*(1-P)) # Nagelkerke scaling factor
  R2 = prs_r2nk*cv*lambda/(1+prs_r2nk*theta*cv*lambda)
return(R2)
}
~~~

